# Path Integration and Spatial Updating Recruit Distinct Cognitive-Neural Mechanisms in Humans

**DOI:** 10.64898/2026.04.11.717901

**Authors:** Xiaoli Chen, Jan Wiener, Mary Hegarty, Thomas Wolbers

## Abstract

Path integration and spatial updating refer to the integration of self-motion information during navigation to update one’s start location and the positions of other locations, respectively. Even though path integration has been described as a fundamental process whose output may serve as a building block for other navigational computations like spatial updating, the exact relationship between path integration and spatial updating is unknown. Here we addressed this question with an eye-tracking behavioral experiment and a subsequent fMRI study. Despite experiencing identical self-motion cues, participants showed differential eye fixation patterns and responded more quickly during spatial updating than during path integration, casting doubt on the fundamental role of path integration. Neuroimaging results showed that the precuneus and the dorsal premotor cortex were more activated during spatial updating, but the precuneus had stronger functional connectivity with the thalamus and the frontal cortex during path integration. Further supporting this dissociation, the two tasks invoked distinct brain-wide inter-regional functional networks. Together, the combined findings of both experiments suggest that spatial updating and path integration are dissociable navigation processes supported by distinct behavioral and neural mechanisms, rather than one process operating on the basis of the other.

## 1. Introduction

Spatial navigation is an essential everyday function that relies on a variety of spatial cues. One important source of spatial information is self-motion cues, which refers to the spatial information derived from internal idiothetic (proprioceptive cues, motor efference copies, vestibular feedback) and external allothetic cues (optic flow) during locomotion. In contrast to static environmental cues (e.g., visual landmarks), navigating with self-motion cues requires continuous integration of these cues over travel distance and direction. Importantly, self-motion-based spatial navigation can be further classified into two different processes: spatial updating and path integration. While spatial updating involves continuously tracking the positions of external objects relative to one’s own moving body, path integration refers to the updating of one’s position relative to a location one has previously occupied (e.g. the origin of travel).

Given that path integration and spatial updating both rely on the integration of self-motion cues and involve online updating of spatial representations, the underlying mechanisms may not be fundamentally different. After all, the home location, just like any other spatial location in the environment, is simply a location external to the navigator’s body (Wang, 2016). Consistent with this view, behavioral studies typically do not draw a strong conceptual distinction between spatial updating and path integration, and the two terms are sometimes even used interchangeably (He et al., 2018; He & McNamara, 2018).

In contrast to this view, neuroscience findings suggest that path integration and spatial updating recruit at least partially distinct brain regions. In human spatial updating, activation of the precuneus is proportional to the number of to-be-tracked objects when the navigator moves in a virtual environment (Jahn et al., 2012; Wolbers et al., 2008), and a virtual lesion to the precuneus induced by repetitive transcranial magnetic stimulation impairs spatial updating performance, demonstrating a causal role of the precuneus (Müller et al., 2018). In contrast, path integration has typically been associated with computations in medial temporal lobe (MTL) structures. For example, hippocampal activation in humans positively predicts individual path integration performance (Wolbers et al., 2007), and reflects the distance from the navigator to the home location (Chrastil et al., 2015). Moreover, grid-cell-like activation in the human entorhinal cortex positively predicts path integration performance in older adults (Stangl et al., 2018). Activation of the human parahippocampal cortex reflects the moment-to-moment changes in the distance to home during path integration (Chrastil et al., 2015). Rodent work provides converging evidence, because both disrupting entorhinal grid cells and hippocampal lesions impair path integration (Gil et al., 2018; Maaswinkel et al., 1999). Beyond the MTL, thalamic head direction cells contribute to the cumulative error inherent in path integration (Valerio & Taube, 2012), highlighting the thalamus as another core component of the path integration circuitry. Taken together, prior findings suggest that whereas spatial updating primarily engages medial parietal regions along the dorsal stream, path integration relies more heavily on MTL structures and the thalamic head-direction network.

Despite extensive investigation of these two processes, it remains unclear whether path integration and spatial updating rely on common or distinct cognitive-neural processes. This uncertainty arises primarily from a lack of direct comparisons between the two processes. To address this gap, we conducted two separate experiments with a novel experimental paradigm that enabled a direct comparison between path integration and spatial updating (Figure 2). In this paradigm, participants moved along a predetermined curved trajectory (the bolded solid curve) while keeping track either of the start position of the trajectory (path integration) or of a target position that was not located on the trajectory (spatial updating). Upon reaching the end of the trajectory, participants had to point to the start position for path integration and to the target position for spatial updating.

Beyond determining whether spatial updating (SU) and path integration (PI) recruit common or differential behavioral and neural mechanisms, we also aimed to more precisely illustrate the relationship between these two processes. Based on the existing literature, we conceived three competing hypotheses. First, the *SU-depends-on-PI hypothesis* argues that path integration is a primitive navigation process (Mittelstaedt & Mittelstaedt, 1980), while more advanced navigation processes, such as spatial updating, build upon it (Wang, 2016; Whitlock et al., 2008). Under this account (Figure 1A), path integration operates in a relatively automatic manner, meaning the start location is continuously updated. Hence, at the time of response, the pointing vector to the start position (e.g., homing vector) is already represented in the cognitive system and only needs to be retrieved. In contrast, because the target location is not automatically tracked during the movement, the pointing vector to the target is not directly available at the time of response. Instead, it must be inferred from the output of the path integrator (i.e., the homing vector). This inference process introduces additional cognitive demands compared to direct retrieval, leading to performance costs like increased reaction time (Klatzky, 1998). Accordingly, this hypothesis predicts worse behavioral performance in path integration than in spatial updating.

**Figure 1.**
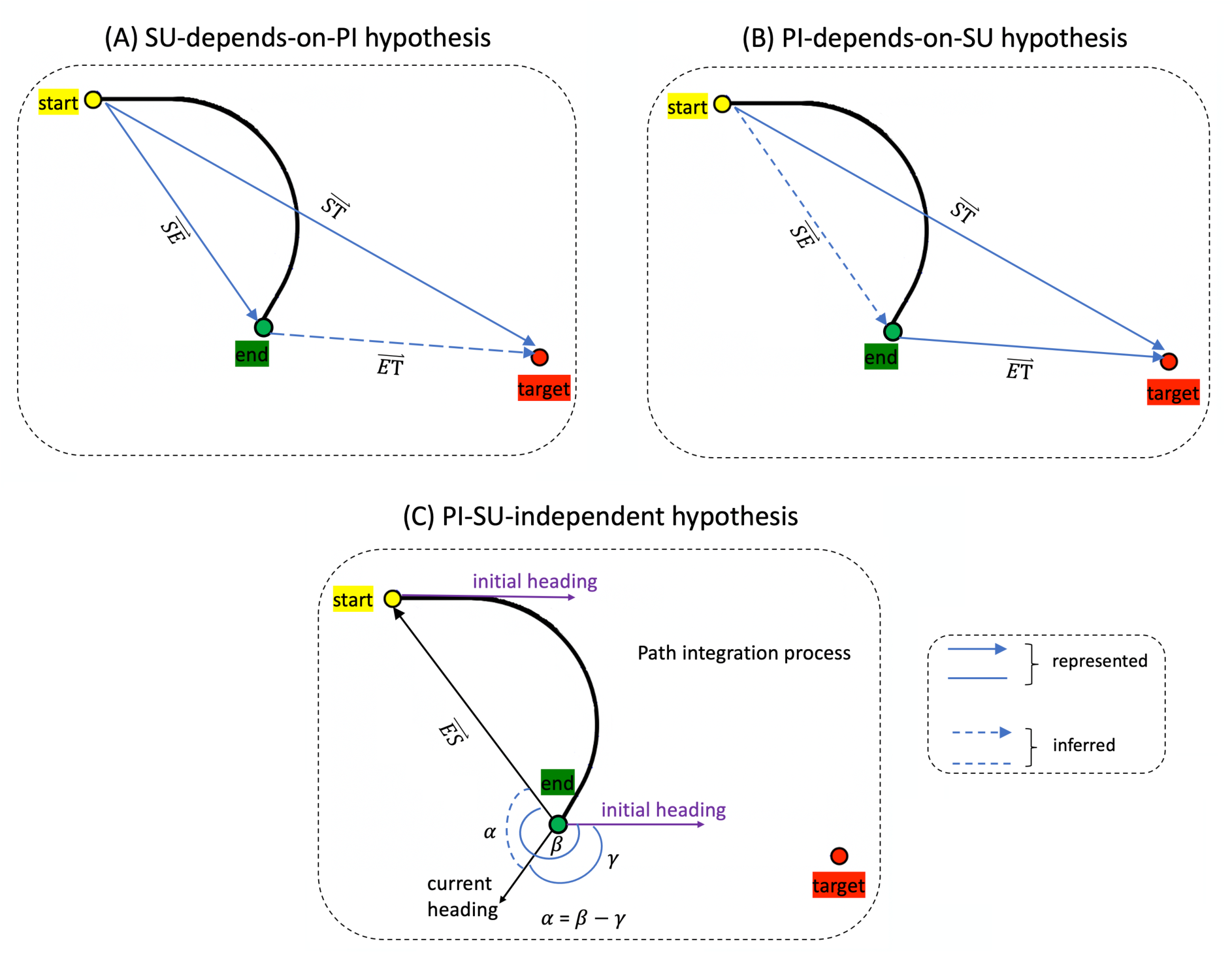
Hypotheses regarding relationships between path integration and spatial updating. In the experimental paradigm, the participant moves along a predetermined trajectory (the bolded solid curve, while keeping track of the start position of the trajectory (path integration) or a target position not on the trajectory (spatial updating). Also refer to Figure 2 for a detailed description of the task. (A) SU-depends-on-PI hypothesis. In path integration, at the end of a trajectory, the home vector 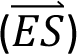 is already represented and only needs to be retrieved. In contrast, in spatial updating, because the target location is not automatically tracked, the pointing vector to the target 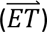 must be inferred from the output of the path integrator (the home vector 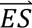) and the stored spatial relation between the start and object positions 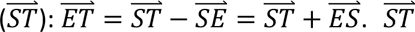 likely have been formed at the beginning of the trial, when the participant views the target, from the starting position. (B) PI-depends-on-SU hypothesis. In spatial updating, at the end of a trajectory, the pointing vector to the target 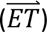 is already represented in the cognitive system and can be directly retrieved. In contrast, in path integration, the home vector 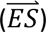 must be computed using the pointing vector and the start-to-target vector (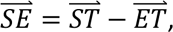 equivalent to 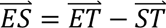) (C) PI-SU-independent hypothesis. In spatial updating, the target location is continuously tracked and updated during movement (same as in Figure 1B). Thus, at the time of response, the pointing vector to the target is directly retrieved. In contrast, in path integration, the pointing direction to home 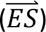 is inferred within a fixed reference frame established by a prominent direction during the task, which corresponds to the initial heading of movement here. Within this reference frame, all directional vectors, such as the current heading (*γ*) and the home direction (*β*), are represented relative to the initial heading of movement. To compute the pointing angle toward home (*α*), the system subtracts the current heading from the home direction (*α* = *β* − *γ*).

In contrast to the SU-depends-on-PI hypothesis, the *PI-depends-on-SU hypothesis* posits that while spatial updating operates automatically, path integration operates on the output of spatial updating (Figure 1B). Under this account, at the end of a trajectory, the pointing vector to the target is already represented in the cognitive system and can be directly retrieved to generate the response. In contrast, the home vector is not tracked during the movement. Instead, it must be computed at the time of response using the pointing vector. This additional computation imposes greater cognitive demands on path integration than on spatial updating. Consequently, this hypothesis predicts better behavioral performance for spatial updating than path integration.

Although the two hypotheses make opposite assumptions, they share one core idea: one process is dependent on the other process, meaning path integration and spatial updating cannot be dissociated. Figure 1C presents a third hypothesis, which posits that the two processes are largely independent (*PI-SU-independent hypothesis*). Like the PI-depends-on-SU hypothesis (Figure 1B), this hypothesis assumes that the target location is continuously tracked and updated relative to participants’ current position during movement. Thus, at the time of response, the pointing vector to the target is readily available, facilitating performance by shortening the reaction time. However, this hypothesis differs from the SU-depends-on-PI hypothesis in its treatment of the pointing-to-home response. Rather than being inferred from the pointing vector to the target, the pointing direction to the start position is inferred within a fixed reference frame established by a prominent direction during the task. The initial heading of movement is a strong candidate for establishing this reference frame, as it remains stable across trials (He et al., 2018). Within this reference frame, all directional vectors, such as the current heading and the home direction, are represented relative to the initial heading of movement. To compute the pointing direction toward home, which is not directly represented, the system subtracts the current heading from the home direction. This additional computation imposes extra cognitive workload, leading to performance cost such as increased reaction time. Because this inference process does not involve the target representation, path integration and spatial updating emerge as largely dissociable processes.

In brief, the SU-depends-on-PI hypothesis predicts better behavioral performance (e.g., faster responses) in path integration than in spatial updating. In contrast, both the PI-depends-on-SU hypothesis and PI-SU-independent hypothesis predict better behavioral performance in spatial updating than in path integration, albeit through different mechanisms. Because the latter two hypotheses shared the same behavioral prediction, fully distinguishing between all three hypotheses requires additional evidence, such as from eye-tracking and neural data.

To preview, our behavioral, eye-tracking, and fMRI results provide converging evidence that spatial updating and path integration are relatively independent processes that recruit dissociable cognitive-neural mechanisms, supporting the PI-SU-independent hypothesis.

## 2. Behavioral Experiment

In the experimental task, participants were briefly presented with an object (the target) in a virtual environment. After memorizing the object’s location, it disappeared, and participants were informed whether the current trial required them to update the object location (spatial updating) or the start location (path integration). Following a subsequent motion phase, during which participants were passively transported along a curved path, they used a joystick to point either to the remembered object or start location. Gaze was recorded binocularly.

### 2.1. Methods

#### 2.1.1. Participants

Twenty naïve healthy participants with normal or corrected to normal vision participated in the experiment (10 women; age range: 20-30 yrs). This sample size was comparable to previous fMRI studies on path integration and spatial updating (Jahn et al., 2012; Wolbers et al., 2007, 2008; Zajac et al., 2019).

Most participants were students from the University of Freiburg and were paid 8€ per hour for participation. Behavioral data for two participants and gaze data of one participant were excluded from the analysis as the corresponding data files were corrupt.

#### 2.1.2. Experimental Stimuli and Paradigm

We used Vizard 2.53 (WorldViz, Santa Barbara, CA) to animate a desktop virtual environment (Figure 2) that participants viewed from a first-person perspective (eye-height: 1.8m). The environment contained a ground floor consisting of white dots that extended 50m in depth and covered the entire field of view laterally (see Figure 2). The remainder of the scene was kept black. To prevent participants from attending to single dots, we used limited lifetime dots. Each dot was moving towards the observer until its initial lifetime (random duration 0 – 2 sec) had expired. It was then redrawn at a random location and its lifetime was reset to 2 sec.

**Figure 2.**
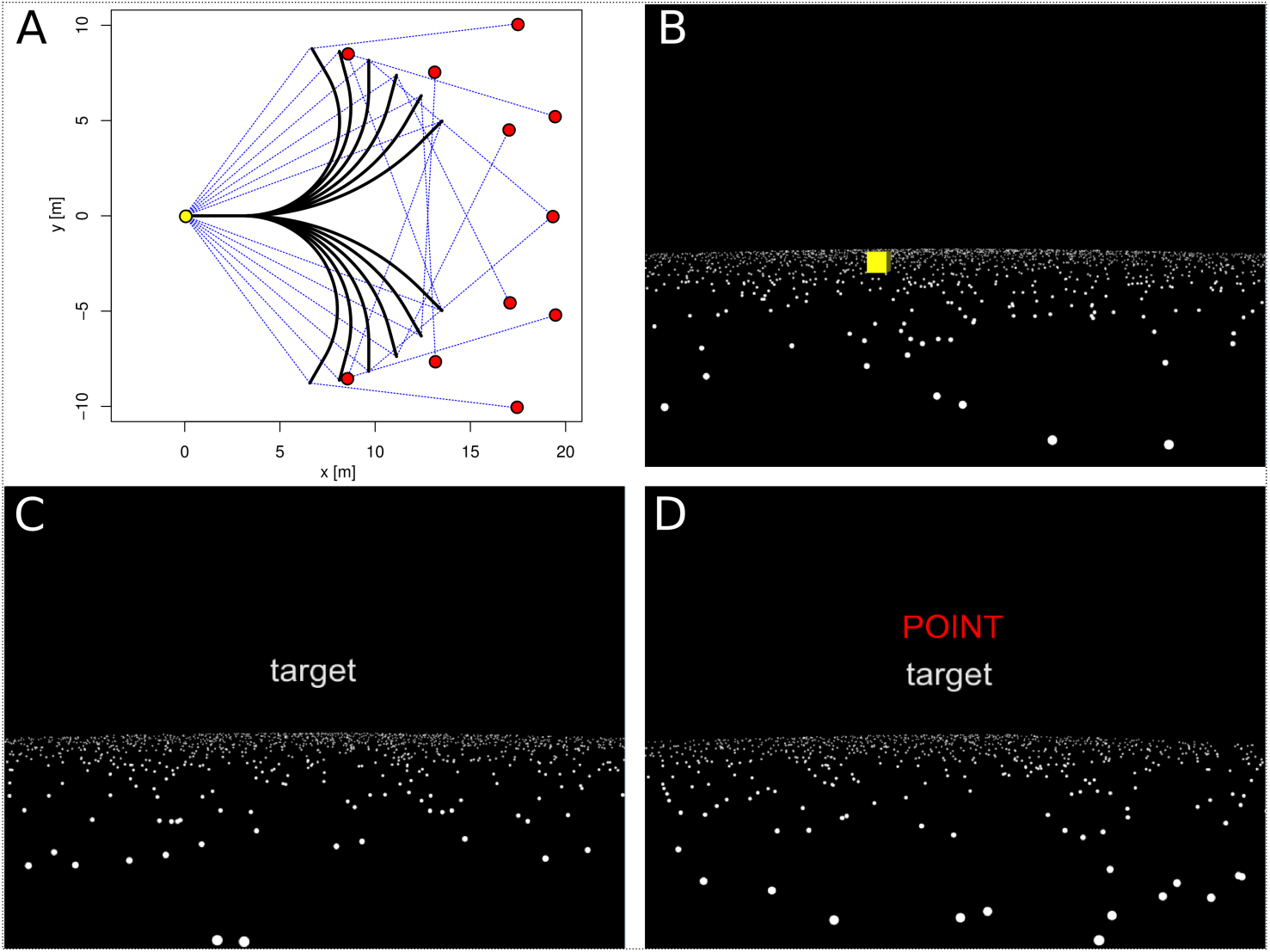
Experimental Setup for the Behavioral Experiment and the FMRI Experiment. **(A)** Paths used in the task. The yellow dot represents the starting position of all the paths (i.e, start). Participants were required to keep track of the start location in path integration trials. The red dots represent to-be-tracked locations (i.e., target) in the spatial updating trials for different paths. Note that for each path, the correct pointing angle to the red dot (target) and the correct pointing angle to the yellow dot (start) are matched in magnitude. **(B-D)** Demonstration of a spatial updating trial. First, a target object (yellow square) appears (B). The target object disappeared once the movement starts (C). After the movement has ended, the participant is instructed to point to the location of the target object (D). Refer to Methods for a detailed description of the task trial.

To study path integration and spatial updating, 12 curved outbound paths were created that were comprised of a short initial straight segment (3m), a curved segment, and a final straight segment (1.5m). The form of the curved segment depended on the overall turning angle along the path. The design ensured identical path length (15m) despite different turning angles (45°, 60°, 75°, 90°, 105°, 120°) between paths (see Figure 2).

For each path one location in space was defined – the object location – at which a geometric object was briefly presented before the onset of the passive transportation. These locations were selected such that the egocentric vector from the final position of the path to the object location was identical to the egocentric vector to the start point of the path, but only differed in the sign. For example, if, after reaching the endpoint of the trajectory, the start location would be at a distance of 7m and 100° to the right, the object location would be at a distance of 7m and 100° to the left (see Figure 2). For training sessions we designed a different set of paths (50°, 65°, 80°,95°, 110°, 125°) that also required different pointing responses.

#### 2.1.3. Apparatus

The virtual environment was displayed at a resolution of 1280 x 1024 pixels on a 20“ CRT monitor. The screen refresh rate was 100 Hz. Participants sat in front of the monitor at a distance of ∼60 cm, such that the resulting visual angle of the stimuli displayed was 37° (horizontally) x 28° (vertically). A standard joystick was mounted onto a desk in from of participants. They responded by deflecting the joystick in direction of the respective target location and confirmed their direction judgments by pressing a joystick button with their index finger. Pointing responses and reaction times were recorded when the participant pressed the button. Eye movements were recorded using a SR Research Ltd. EyeLink II eye tracker, sampling pupil position at 500 Hz. The Eyelink II eye-tracker is a head-mounted system that does not require the participants’ head to be constrained. The eye-tracker was calibrated using a 9-point grid. A second 9-point grid was used to calculate the accuracy of the calibration.

#### 2.1.4. Procedure

All participants completed one training session and one experimental session. Each trial started with a static presentation of the virtual environment lasting 7.75 sec. During the first 5 seconds – the encoding phase –, participants saw one geometrical object (cylinder, cube, pyramid or sphere) positioned at one out of 12 possible positions on a ground plane. The ground plane consisted of 3000 white dots scattered randomly across the surface (Figure 2).

Participants were instructed to memorize the locations as precisely as possible during this encoding phase. In order to prevent any interference effects from previous trials, the shape of the object was changed from trial to trial to. Next, the object sank into the ground and disappeared (duration: 2 sec), and the animation then remained still for another 0.75 sec. Participants were then instructed via a text message on the screen whether they had to point to the object location (*spatial updating* trial) or to the start location of the travel (*path integration* trial) upon reaching the end of the path. They were instructed to respond as quickly and as accurately as possible. During the subsequent movement phase (duration: 12.5 sec), participants were passively transported along the current path. Given that self-motion perception in virtual environments is most accurate when displacement velocity resembles natural locomotion (Ellmore & McNaughton, 2004), we adopted a speed of a moderately paced walk (1.5m/sec) and set eyeheight at 1.5m.

In the *training phase*, participants were presented with a number of paths and were asked to point back to the starting place or to an object. Training paths had different lengths and turning angles than test paths. Participants were presented with a random selection of training trials until they understood the experimental procedure. During the *test phase*, participants were passively transported along all 12 test paths twice (see Figure 2); each path was once paired with the path integration task and once with the spatial updating task. The order randomized of the presentation of the 24 trials was randomized.

#### 2.1.5. Statistical analysis of behavioral data

For each trial, we measured absolute pointing error and reaction time. Larger pointing error for low vs. high dot density would reveal uncertainty related to the amount of self-motion information provided by the ground plane. In addition, differences between start and target trials would be indicative of different processing demands related to updating locations occupied by the observer or by the object at encoding. To assess the effects of our experimental manipulations on both dependent variables, we performed two-way repeated measures analyses of variance (ANOVA) with task and dot density as within-group factors. Violations of sphericity were addressed with Greenhouse-Geisser correction of the degrees of freedom.

### 2.2. Results

#### 2.2.1. Pointing Performance

Pointing accuracy was measured as absolute pointing error, which equaled to the absolute difference between the correct pointing angle and the actual pointing angle. Absolute pointing error did not differ between left and right turn paths (t(17)= -0.56, p = 0.58) and was therefore pooled for the remaining analyses. As shown in Figure 3a, absolute pointing error increased with increasing turning angle, but it did not differ between path integration and spatial updating trials. A 2x6 repeated measures ANOVA revealed a significant main effect of absolute turning angle (F(5, 85) = 3.49, p < 0.01), but neither a main effect of task (F(1, 17) = 0.01, p = 0.94) nor a significant interaction (F(5, 85) = 0.71, p = 0.61).

**Figure 3.**
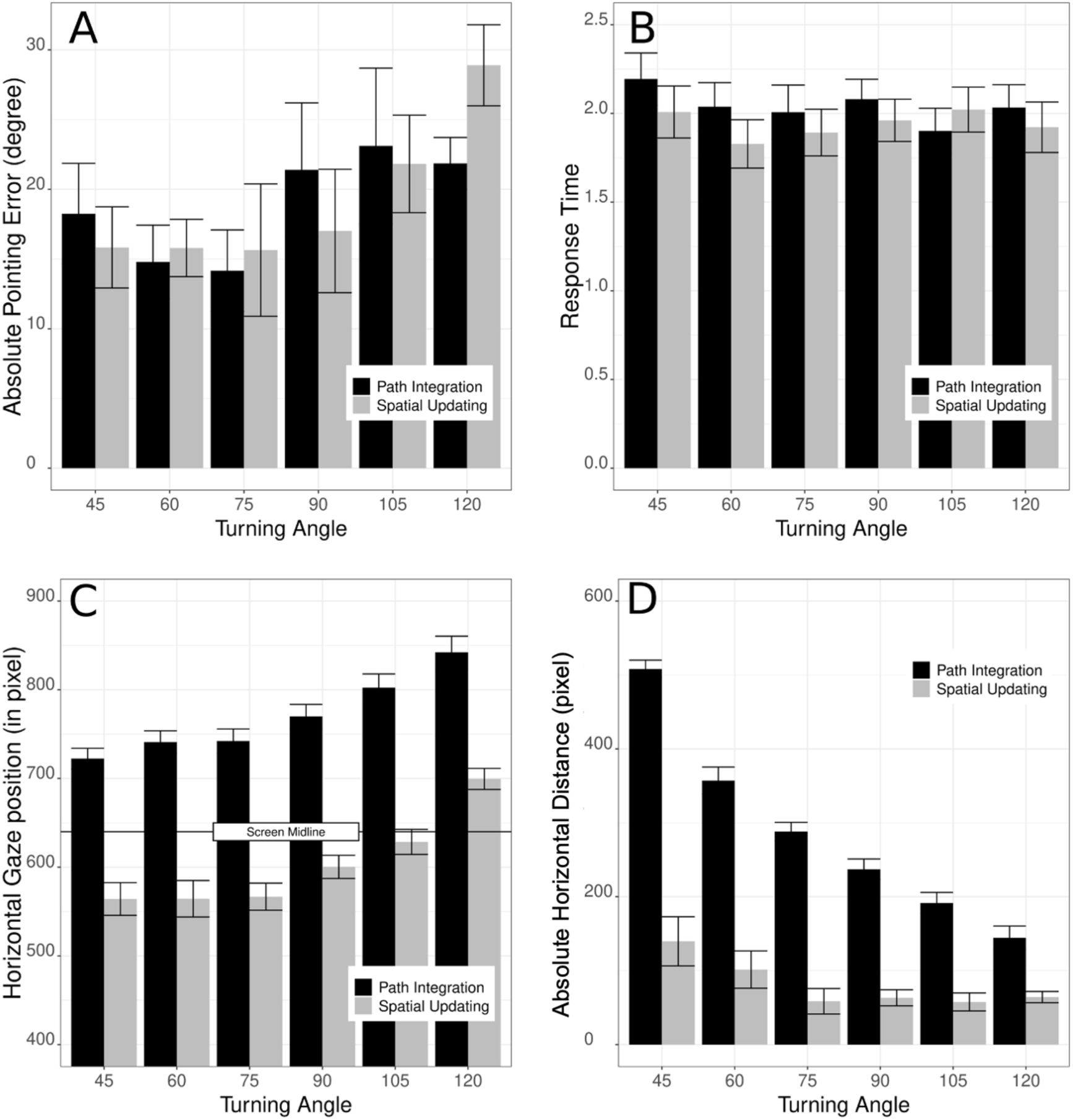
Behavioral Performance and Eye-Tracking Results in the Behavioral Experiment. (A) Absolute pointing error increased as a function of turning angle during the outbound path. (B) Average response time was shorter for spatial updating than for path integration. (C) Average horizontal gaze position as a function of turning angle. (D) Absolute horizontal distance between gaze and object location. In all plots, bar height represents the mean and error bars represent ±SEM.

In contrast, response time data showed a slightly different pattern (Figure 3B): participants responded faster during spatial updating (mean: 1936 msec) than during path integration (mean: 2045 msec) trials (main effect of task: F(1, 17) = 5.74, p = 0.03), indicating a large effect. Reaction times were independent of absolute turning angle (F(5, 85) = 1.31, p = 0.27), and there was no interaction between both factors (F(5, 85) = 1.06, p = 0.39).

Taken together, these results demonstrate that representations of locations became less accurate with larger turning angles, but this effect was similar for both tasks. Despite this comparable task difficulty, participants responded consistently faster when the task required the updating of an external location. This result is difficult to reconcile with the SU-depends-on-PI hypothesis (Figure 1A), which suggests that path integration is a fundamental process underlying various forms of updating spatial locations. In contrast, this result is consistent with both the PI-depends-on-SU hypothesis (Figure 1B), which proposes that path integration operates on the output of spatial updating, and the PI-SU-independent hypothesis (Figure 1C), which proposes that path integration and spatial updating rely on largely dissociable mechanisms.

#### 2.2.2. Eyetracking Results

To analyze the eye-tracking data, we first pooled data for paths turning left and turning right by mirroring the gaze data for the horizontal position for left turn paths. We then synchronized data between trials at the start of the translation phase, which allowed for comparisons of gaze behavior during the translation phase between tasks and different path configurations. Gaze data for the horizontal and vertical screen position were analyzed independently.

Average vertical gaze position did not differ between the two tasks (main effect of task: F(1,18) = 0.06, p = 0.81), nor was it affected by the varying turning angles (main effect of turning angle: F(5,90) = 0.89, p = 0.49, interaction: F(5, 90) = 0.69, p = 0.63). In contrast, a repeated measures ANOVA on horizontal gaze position (Figure 3C) revealed significant main effects of task (F(1, 18) = 113.92, p < 0.001) and turning angle (F(5, 90) = 38.03, p < 0.001), but no significant interaction (F(5, 90) = 0.55, p = 0.74). Specifically, gaze was directed to the right hemifield of the screen during path integration trials, while it was directed to the hemifield in which the object had initially been shown in all spatial updating trials (except for the path with a 120° turning angle). Moreover, the deviation from straight ahead increased as the turning angle of the outbound path increased.

To better understand the nature of these effects, we compared participants’ mean gaze position during the different outbound paths to the location where the object had been presented (note that the object was no longer visible, as it had already disappeared before motion onset). As apparent in Figure 4, participants closely followed the remembered object location with their gaze until it “left” the screen during spatial updating trials. This gaze following behavior was completely absent during path integration trials. We confirmed this impression with an analysis in which we calculated the absolute horizontal distance (in pixel) between the participants’ gaze and the object location. This analysis was restricted to the part of the translation during which the object location was still on the screen. A one way ANOVA revealed a significant main effect of task (F(1, 18) = 155.78, p < 0.001), demonstrating that the average distance between gaze and object location on the screen was significantly smaller for spatial updating trials (109 pixel) than for path integration trials (332 pixel; Figure 3D).

**Figure 4.**
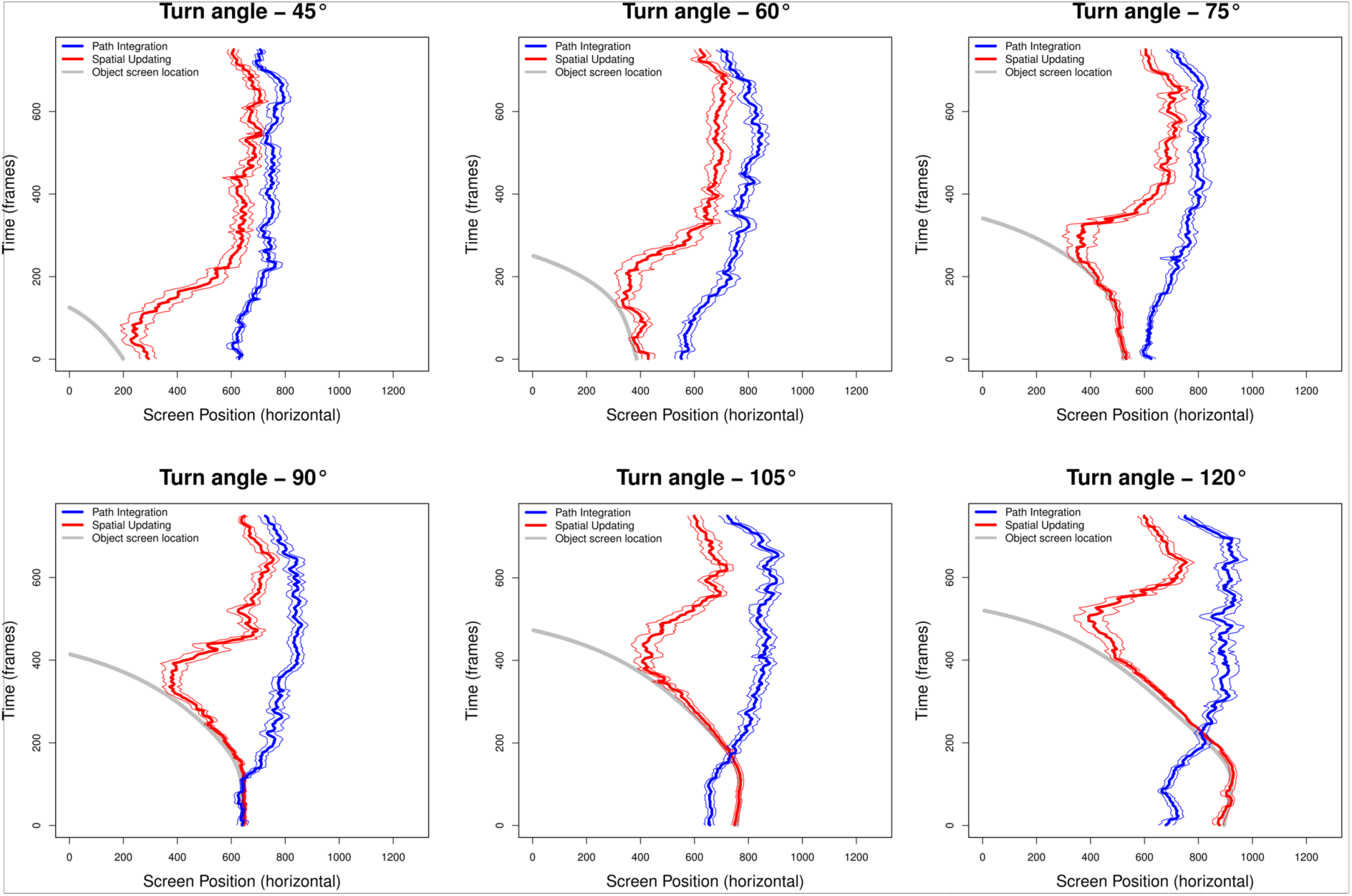
Gaze Position in the Behavioral Experiment. Average horizontal gaze position during the motion phase for spatial updating (red) and path integration (blue) trials for the different paths. The screen position of where the target object had initially been presented is indicated by the grey line.

Figure 4 also reveals that the difference in horizontal gaze position between path integration and spatial updating trials was markedly reduced once the remembered object location left the screen. Specifically, participants moved their fixation back in the direction of motion in spatial updating trials, presumably to bring the focus of expansion – an area of minimal optic flow with maximum information content for judging self-motion direction on curvilinear paths – into their focus of attention (Authié & Mestre, 2012).

In brief, the eye tracking results revealed distinct gaze patterns between spatial updating and path integration. During spatial updating, participants’ gaze continuously tracked the target location until it moved out of the screen. In contrast, during path integration, they primarily fixed their gaze in the direction of self-movement. These results contradict the PI-depends-on-SU hypothesis. Under this hypothesis, the output of spatial updating is required to compute the homing vector in path integration; therefore, it predicts that participants in the path integration task should exhibit a eye-gaze pattern similar to those observed in spatial updating trials, which was not supported by the data.

In addition, the results also conflict with the SU-depends-on-PI hypothesis, which posits that spatial updating is not automatic but instead dependent on the output of path integration. This claim is inconsistent with the finding that participants’ gaze continuously followed the target location during spatial updating, indicating automatic updating of the target’s position relative to the self. Notably, this eye-tracking finding of automatic updating may help explain the earlier behavioral finding that participants responded faster during spatial updating than during path integration.

In contrast, the eye-tracking findings are broadly consistent with the PI-SU-independent hypothesis, as the distinct eye-movement patterns observed across the two tasks suggest the engagement of separate underlying mechanisms.

#### 2.2.3. Summary of Behavioral and Eyetracking Results

Two important conclusions can be drawn from the behavioral experiment. First, while both tasks were of equal difficulty, participants were quicker to respond in spatial updating trials. This behavioral finding is inconsistent with the SU-depends-on-PI hypothesis, but is consistent with both the PI-depends-on-SU hypothesis and the PI-SU-independent hypothesis.

Second, during spatial updating, participants initially tracked the remembered location of an external object by following it with their eyes. However, once this position left the field of view, participants directed their attention in the direction of motion. In contrast, during path integration, this latter strategy was applied from the beginning of a trial, suggesting that participants focused on the area of minimal optical flow, which maximizes the ability to monitor the direction of self-motion as accurately as possible (Authié & Mestre, 2012). This eye-tracking finding is inconsistent with the PI-depends-on-SU hypothesis and the SU-depends-on-PI hypothesis, but is broadly consistent with the PI-SU-independent hypothesis.

## 3. FMRI Experiment

Considering both the behavioral and eye-tracking results, the PI-SU-independent hypothesis appears the most plausible. However, direct evidence for this account remains limited, as behavioral and eye-tracking measures provide only indirect insight into the specific cognitive processes underlying both tasks. To more directly investigate the relationship between path integration and spatial updating – and to better distinguish among the three competing hypotheses – we conducted an fMRI experiment to examine the underlying neural mechanisms.

In the fMRI experiment, participants performed the same spatial updating and path integration tasks as in the behavioral experiment. In addition, we introduced a perceptual manipulation by varying the number of dots on the ground plane (750 vs. 3000). Critically, participants were required to continuously fixate a central cross in both conditions to ensure that potential differences in behavioral performance or BOLD activation patterns could not be driven by differences in eye movement patterns.

### 3.1. Methods

#### 3.1.1. Participants

Sixteen healthy volunteers with normal or corrected-to-normal vision gave written informed consent to participate in this study, which was conducted at UC Santa Barbara’s Brain Imaging Center and was approved by the local ethics committee. All participants understood the instructions without difficulties and none were aware of the hypotheses at the time of testing.

The sample size was similar to previous fMRI studies on path integration and spatial updating (Jahn et al., 2012; Wolbers et al., 2007, 2008; Zajac et al., 2019).

#### 3.1.2. Experimental Stimuli

The paradigm used in the fMRI experiment was very similar to the behavioral experiment; however, two key differences were introduced: participants were required to fixate during self-motion to eliminate eye movement confounds, and we introduced a perceptual manipulation in terms of varying the number of dots. Hence, a 2 x 2 factorial design with factors *task* (path integration vs. spatial updating) and *dot density* (750 vs. 3000) served to characterize the neural mechanisms that differentiate path integration from spatial updating. As in the behavioral experiment, trials consisted of three phases: an initial encoding phase followed by a movement phase and a subsequent retrieval phase. Each trial started with a static presentation of the virtual environment lasting 7.75 sec. During the first 5 seconds – the encoding phase –, participants saw one geometrical object (cylinder, cube, pyramid or sphere) positioned at one out of 12 possible positions on a ground plane. The ground plane consisted of 750 or 3000 white dots scattered randomly across the surface (Figure 2). Participants were instructed to memorize the location of the object as precisely as possible. Importantly, the shape of the object was changed from trial to trial to prevent any interference effects from previous trials. Next, the object gradually sank into the ground until complete disappearance (duration: 2 sec), and the animation then remained still for another 0.75 sec. Finally, a central white fixation cross and the task instruction for the current trial were displayed (Figure 2): when the word ‘start’ appeared above the fixation cross, participants had to point towards the starting location at the end of the trial. Conversely, the word ‘target’ signaled that participants would have to point to the remembered location of the target.

In the subsequent movement phase (duration: 12.5 sec), participants experienced a passive translation along one of eight curved paths, using the same layouts as in the behavioral experiment (Figure 2). Participants were instructed to pay close attention to the optic flow provided by the limited lifetime dots – while maintaining fixation on the central cross – in order to monitor their movements through space. In the final retrieval phase, the fixation cross turned red to signal to participants that they were required to point towards either the starting location (start trials) or to the object’s remembered location (target trials) within a 3.5 sec interval. Participants indicated their response with an MR-compatible joystick (Mag Design and Engineering, Sunnyvale, CA) that was positioned on the right thigh. Participants deflected the joystick with the right hand and pushed a button with the left hand when the joystick was oriented in the desired direction. Pointing responses and reaction times were recorded when the participant pressed the button. Participants were not given any feedback about their performance, and a black screen with the white fixation cross was presented during inter-trial intervals that randomly varied between 3 and 5 sec.

#### 3.1.3. Procedure

All participants completed two training sessions followed by one experimental session. We first performed extensive training to eliminate learning and habituation effects. Detailed instructions about both tasks were followed by a real-world version outside the MR environment: after encoding the position of the experimenter, participants walked along a straight path for a randomly chosen distance without turning their heads. When the experimenter was out of sight (i.e. behind the participant), participants were asked to stop and point either towards the remembered position of the experimenter (*spatial updating* trial) or towards their starting location (*path integration* trial). Participants were not allowed to turn their heads and gave the responses with the joystick that was also used for the virtual environments. Next, participants completed ten training trials presented on a standard computer monitor. During these training trials, pointing responses were followed by instant feedback: an arrow was presented parallel to the ground plane that indicated the correct pointing direction. This procedure proved successful since all participants were subsequently able to perform the task with relatively low pointing errors (see Results). After being positioned within the bore of the magnet, participants were given another set of ten training trials without concurrent fMRI recording to familiarize them with performing the task while in a horizontal position. In the subsequent experimental session, 24 target and 24 start trials were presented in a pseudo-randomized order. Half of these trials showed a low density ground plane (750 dots), the other half showed a high density ground plane (3000 dots).

During encoding, the object could only appear at one of 12 possible positions, six in the right and six in the left hemifield. The same positions were used for each of the four trial types (start/target * high/low dot density). Participants were not informed that the target objects were presented at the same 12 positions in each condition, and none of the experimental trials provided behavioral feedback. All trials were scanned in one fMRI session. After the experiment, participants were debriefed with regard to the replication of the target locations, but none of them reported noticing any patterns in the stimulus positions.

In sum, our paradigm ensured that during the initial encoding phase of each trial, participants had to encode the location of the object, given that they did not know whether they would have to point to the object or to the starting location at retrieval. During the movement phase, participants only experienced optic flow and had to use these self motion cues to update either the object or the starting coordinates. The varying number of dots on the ground plane provided a variation of perceptual stimulation in both conditions.

#### 3.1.4. MRI Acquisition and Imaging Processing

MR scanning was performed on a 3T MRI Scanner (Siemens Trio) with a standard headcoil. Thirty-six axial slices (slice thickness: 3mm, gap: 0.5mm) were acquired using a gradient echo echo-planar T2*-sensitive sequence (TR=1.92 sec, TE=30 ms, flip angle 75°, matrix 64*64, field of view 192*192 mm). An LCD projector back-projected the virtual environment on a screen positioned behind the head coil. Participants lay on their backs within the bore of the magnet and viewed the stimuli binocularly via a 45° mirror that reflected the images displayed on the screen. To minimize head movement, all participants were stabilized with tightly packed foam padding surrounding the head.

Image processing and statistical analysis were carried out using SPM12. All volumes were realigned to the first volume, spatially normalized to an EPI template in a standard coordinate system (Evans et al., 1993) and finally smoothed using a 6 mm full-width at half-maximum isotropic Gaussian kernel.

#### 3.1.5. Statistical Analysis of FMRI Data

##### 3.1.5.1. First-level GLM to detect brain activation differences between tasks

To detect brain activation differences between path integration and spatial updating, we constructed a first-level GLM.

At the single-participant level, we applied a high-pass filter to remove low frequency artifacts. Design matrices containing separate regressors for each of the four trial types (spatial updating / path integration * low / high dot density) were specified. We also included six regressors coding for rotation and translation of the head as obtained by the realignment procedure. Encoding, movement and retrieval intervals were modeled as boxcar functions convolved with a hemodynamic response function (HRF); however, trials in which participants failed to respond within the 3.5 sec interval were modeled as separate regressors and were discarded during statistical analyses. Specific effects were tested with appropriate linear contrasts of the parameter estimates, and the corresponding contrast images were entered into a random effects analysis.

##### 3.1.5.2. Psychophysiological Interaction (PPI) Analysis

The abovementioned brain activation analysis could identify brain regions whose activation differed between path integration and spatial updating. To investigate whether these brain regions also exhibited differential connectivity pattern with other brain regions between these two task types, we conducted a PPI analysis.

PPI is a connectivity analysis to identify regions whose activity depends on the interaction of a psychological factor and a physiological factor. To investigate brain regions whose coupling with the precuneus differed between spatial updating and path integration, we specified task type (path integration vs. spatial updating) as the psychological factor and the time course of ‘neural activity’ in the precuneus as the physiological factor. The precuneus was defined as the cluster of significant voxels identified in our previous study (Wolbers et al., 2008) as involved in spatial updating. First, the time course of BOLD response of the precuneus was extracted, which was then deconvolved to obtain the underlying neural activity. Second, we constructed the first-level participant-wise GLM, which comprised a main effect of the psychological factor (i.e., task type), a main effect of the physiological factor (i.e., deconvolved time course of the BOLD response in the precuneus), and their interaction. The interaction term indicates whether the effective connectivity from the precuneus to a certain brain region differs between tasks. The PPI analyses were implemented in SPM12, following the example documented in the SPM12 manual (Chapter 36 – Psychophysiological Interactions). For the group-level analysis, we focused on the interaction effect and conducted a nonparametric permutation test with multiple comparisons correction across the entire search volume; we adopted the cluster-level inference, with voxel-wise t value > 3 (Nichols & Holmes, 2002).

##### 3.1.5.3. Multi-Edge Pattern Analysis of Whole-Brain Inter-Region Functional Connectivity

The abovementioned PPI analysis could identify a seed region’s connectivity differences between tasks. To investigate the brain-wide connectivity pattern differences between path integration and path integration, we conducted the multi-edge pattern analysis at the entire brain level.

To construct the population-level functional connectivity matrix, first, we obtained the 264 representative brain nodes from the paper by Power et al. (2011). Second, we extracted the time series for each node corresponding to the two different task types, based on the same first-level GLM used for the preceding PPI analysis. Third, for each task type, we constructed the inter-region functional connectivity matrix by calculating the pairwise Pearson correlations among the 264 nodes. Forth, we constructed the functional connectivity matrix across all the participants: each participant had two rows corresponding to path integration and spatial updating respectively, resulting in 16 * 2 = 32 rows in total; the total number of columns corresponds to the number of elements in calculating pairwise correlations among the 264 Power nodes (= (264*264-264)/2 = 34716 elements).

After obtaining the population-level functional connectivity matrix, first, we performed the principal-component-analysis (PCA) to extract 29 principal components, which together explained 96.63% of the total variance. Second, we performed leave-one-participant-out cross-validation classification analysis, using a random forest classifier from the scikit-learn toolbox as implemented in Python 3.8. In other words, in each round, two rows belonging to the same participant were used as the test data in each round, whereas the data from all the other participants were used as the training data. The actual classification accuracy was calculated based on the observed data. Third, to obtain the surrogate distribution of classification accuracy, in each permutation, we shuffled the two different task labels (start vs. target) within each participant and calculated classification accuracy based on the shuffled data. This procedure was repeated for 5000 times, resulting in the surrogate distribution of classification accuracy. Finally, we compared the actual classification accuracy to the surrogate distribution and calculated the proportion of values in the distribution being greater than the actual classification accuracy (i.e., p_1_tailed_).

##### 3.1.5.4. Functional localizer of human motion complex (hMT+)

The human hMT+ is along the visual motion and navigation pathways. It is well-documented that this region plays a critical role in the processing of optic flow information (Duffy & Wurtz, 1991). Therefore, we investigated how this region’s activation was affected by task type and dot density manipulation. A separate fMRI paradigm served to functionally localize hMT+ in each participant, according to previously established procedures (Huk et al., 2002; Morrone et al., 2000). Briefly, a stimulus of alternating moving and stationary dot patterns was presented in a circular aperture with interleaved rest periods. Moving dots (velocity: 17.1°/s) traveled toward and away from the fixation cross for 16 sec, followed by a 16-s stationary dot field, and a 20-s blank screen. Participants fixated at all times. Regions showing greater BOLD responses to moving than stationary dots were then identified using a first-level GLM and used as functional regions of interest for subsequent analyses.

### 3.2. Behavioral Results: Pointing Performance

As in the behavioral experiment, pointing accuracy was calculated as absolute pointing error, i.e., the absolute angular difference between the correct pointing direction and the participant’s actual pointing direction. As shown in Figure 5A, absolute pointing error did not differ between left and right turn paths (t(15)= -0.21, p = 0.84) and was therefore pooled for the remaining analyses. A repeated-measures ANOVA with task type (path integration vs. spatial updating), dot density (high vs. low), and turning angle (from 45 to 120 in step of 15, six levels) as independent variables only revealed a significant main effect of turning angle, F(5,75) = 8.92, p < 0.001, η_p_^2^ = 0.37), because absolute pointing error increased with increasing turning angles. All remaining effects were insignificant (Fs < 2.04, ps > 0.12). In particular, the main effect of task was insignificant (F(1,15) = 1.29, p = 0.27, η_p_^2^= 0.08), neither were the interaction effects involving task type (Fs < 2.04, ps > 0.12). These results indicate that path integration and spatial updating tasks did not differ in overall task difficulty.

**Figure 5:**
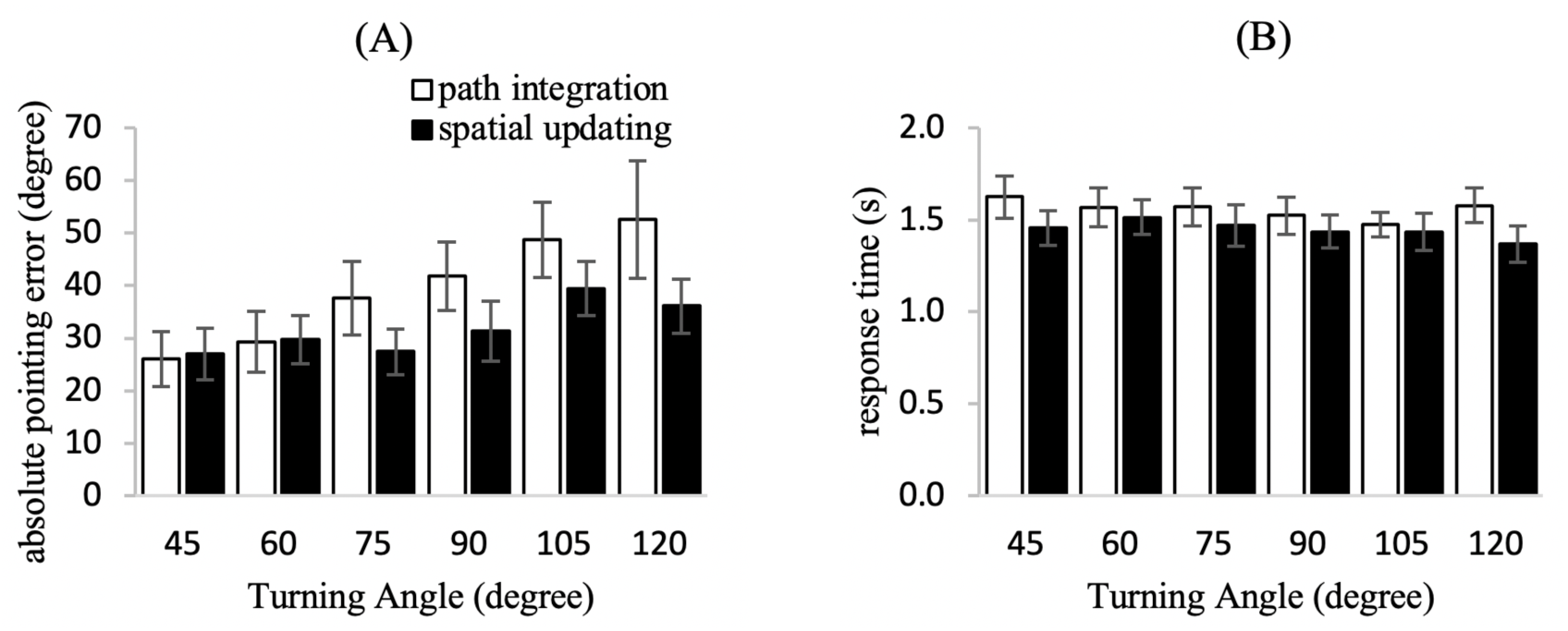
Behavioral Results of the FMRI experiment. (A) Absolute pointing error is plotted as a function of task type and turning angle. The main effect of turning angle was significant, meaning that absolute pointing error increased as turning angle increased, regardless of task type. (B) Response time is plotted as a function of task type and turning angle. The main effect of task type was significant, meaning that response time was longer in path integration than spatial updating, regardless of turning angle. In both plots, bar height represents the mean and error bars represent ±SEM.

In a corresponding ANOVA on response times (Figure 5B), with task type, dot density, and turning angle as independent variables, the only significant effect was the main effect of task type (F(1,15) = 13.82, p = 0.002, η_p_^2^ = 0.48, Cohen’s f = 0.96), indicating a large effect. Specifically, participants performed the task more quickly in spatial updating trials (mean: 1446 msec) than in path integration trials (mean: 1557 msec). None of the other effects were significant (Fs < 3.50, ps > 0.08).

To check for systematic changes in pointing performance over the course of the experiment, we computed the correlation between absolute pointing error and trial number for each participant and each of the four conditions. We then transformed those correlations to normally distributed variables using Fisher’s z-transformation, averaged the z-values across participants and transformed the resulting means back to correlation coefficients. The correlation was not significant in any conditions, and less than 1% of the performance variability was related to increasing familiarity with the tasks both for spatial updating (r = 0.066; high dot density, t(15) = 1.51, p = 0.15; low dot density, t(15) =-0.29, p = 0.78) and for path integration trials (r= -0.01; high dot density, t(15) = -0.09, p = 0.93; low dot density, t(15) = -0.11, p = 0.92). These results indicate that the extensive pre-training (see Methods) successfully prevented learning and habituation effects from occurring over the course of the experiment.

Altogether, while increasing the number of dots did not have an effect on pointing accuracy or reaction time, participants responded significantly faster when the task called for updating the position of an external object. Importantly, pointing accuracy did not differ between both tasks, which demonstrates that the overall task difficulty was comparable. These results replicate the effects observed in the behavioral experiment, indicating that equivalent processes were recruited across different experimental settings.

In brief, the fMRI experiment replicates the previous behavioral experiment. First, pointing accuracy did not differ between path integration and spatial updating, indicating comparable task difficulty. Second, faster responses for spatial updating than path integration again contradict the SU-depends-on-PI hypothesis. However, as mentioned earlier, these behavioral findings cannot distinguish between the PI-depends-on-SU hypothesis and the PI-SU-independent hypothesis, necessitating an examination of the neural data.

### 3.3. FMRI results

#### 3.3.1. Univariate Analysis of Task and Dot-Density Effects

As our 2 x 2 design provided a sensory manipulation (dot density) and a contextual manipulation (task), we first tested for the main effects of task. We applied the cluster-extent threshold technique in SPM12, with a cluster-defining-threshold at p < 0.001 (T > 3.74). This technique appropriately controls the family-wise error at p < 0.05 (Eklund et al., 2016). We later confirmed the results with a nonparametric permutation test with the same cluster-defining threshold (Nichols & Holmes, 2002).

Path integration trials did not evoke stronger BOLD responses than spatial updating trials anywhere in the brain. In contrast, spatial updating trials produced significantly greater activation in a large cluster in the right precuneus and in a second large cluster labeled as the middle frontal gyrus by the AAL atlas, which anatomically corresponds to the dorsal premotor cortex (PMd) (Figure 6, Table 1). This pattern replicates our previous findings demonstrating a central role of the precuneus and the dorsal premotor motor in spatial updating (Wolbers et al., 2008). In addition, as shown in the right panels of Figure 6, activation in these regions was unaffected by dot density, indicating insensitivity to low-level sensory differences when engaged in navigation. Both clusters also reached statistical significance when the nonparametric permutation test was applied (precuneus cluster, p_FWE-corr_ = 0.0064; dorsal premotor cortex cluster, p_FWE-corr_ = 0.0138).

**Figure 6:**
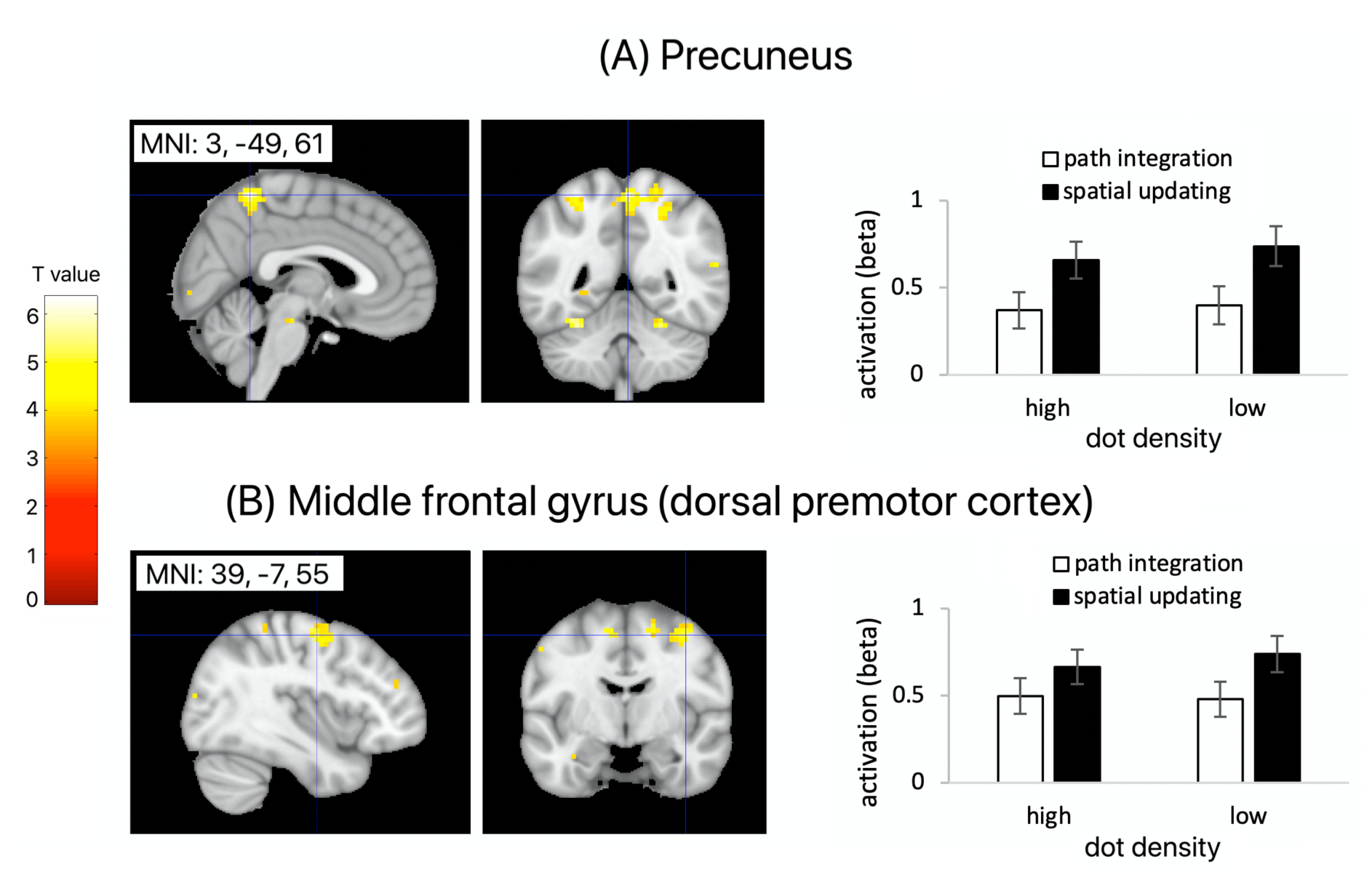
Main Effect of Task Type on BOLD Responses in the Univariate FMRI Analysis. **(A)** Precuneus was more strongly activated in spatial updating trials than path integration trials, regardless of dot density. **(B)** Middle frontal gyrus (dorsal premotor cortex) was more strongly activated in spatial updating trials than path integration trials, regardless of dot density. Results are displayed on the MNI template at a threshold of p_uncorrected_ < 0.001. In the right panels, we extracted mean beta estimates (in arbitrary units) for the significant clusters, which are plotted as a function of task type and dot density. Error bars represent SE of the mean.

**Table 1:** Main Effect of Task Type in the Univariate FMRI Analysis. Using the parametric cluster-based analysis (voxel defining threshold p_uncorrected_ < 0.001), we observed two significant clusters with stronger activation in spatial updating than path integration trials. Cluster size was defined with voxel-wise p_uncorrected_ < 0.001. RH/LH – right/left hemisphere.

| Cluster | Region | Coordinate<br>(x, y, z, in mm) | | Voxel-level<br>(t-score) | cluster size<br>(k) | $p_{\text{FWE-corr,cluster-based}}$ |
| --- | --- | --- | --- | --- | --- | --- |
|  |  | LH | RH |  |  |  |
| Cluster 1 | Precuneus |  | 3, -49, 61 | 6.35 | 142 | < 0.001 |
|  | Inferior parietal cortex |  | 27, -52, 52 | 5.13 |  |  |
|  | Superior parietal cortex |  | 21, -52, 61 | 5.03 |  |  |
| Cluster 2 | Middle frontal gyrus |  | 39, -7, 55 | 5.11 | 87 | 0.003 |
|  | Superior frontal gyrus |  | 18, -7, 61 | 4.97 |  |  |
|  | Precentral gyrus |  | 27, -1, 52 | 4.53 |  |  |

We also tested for the effect of dot density on activation during the motion phase. As shown in Figure 7 and Table 2, this analysis revealed stronger BOLD responses in large cluster of voxels in early visual areas (incl. V1 and V2; cluster size = 544 at p_uncorrected_ < 0.001; Figure 7A) when more dots were present. This effect that was independent of the task being carried out. We also observed a similar effect in a small cluster of voxels encompassing the right hippocampus and thalamus (cluster size = 52 at p_uncorrected_ < 0.001; Figure 7B). These two clusters also reached statistical significance when a nonparametric permutation test was applied (calcarine cluster, p_FWE-corr_ = 0.0002; hippocampus cluster, p_FWE-corr_ = 0.0256). In contrast, the reverse contrast (low > high dot density) did not yield any significant effects.

**Figure 7:**
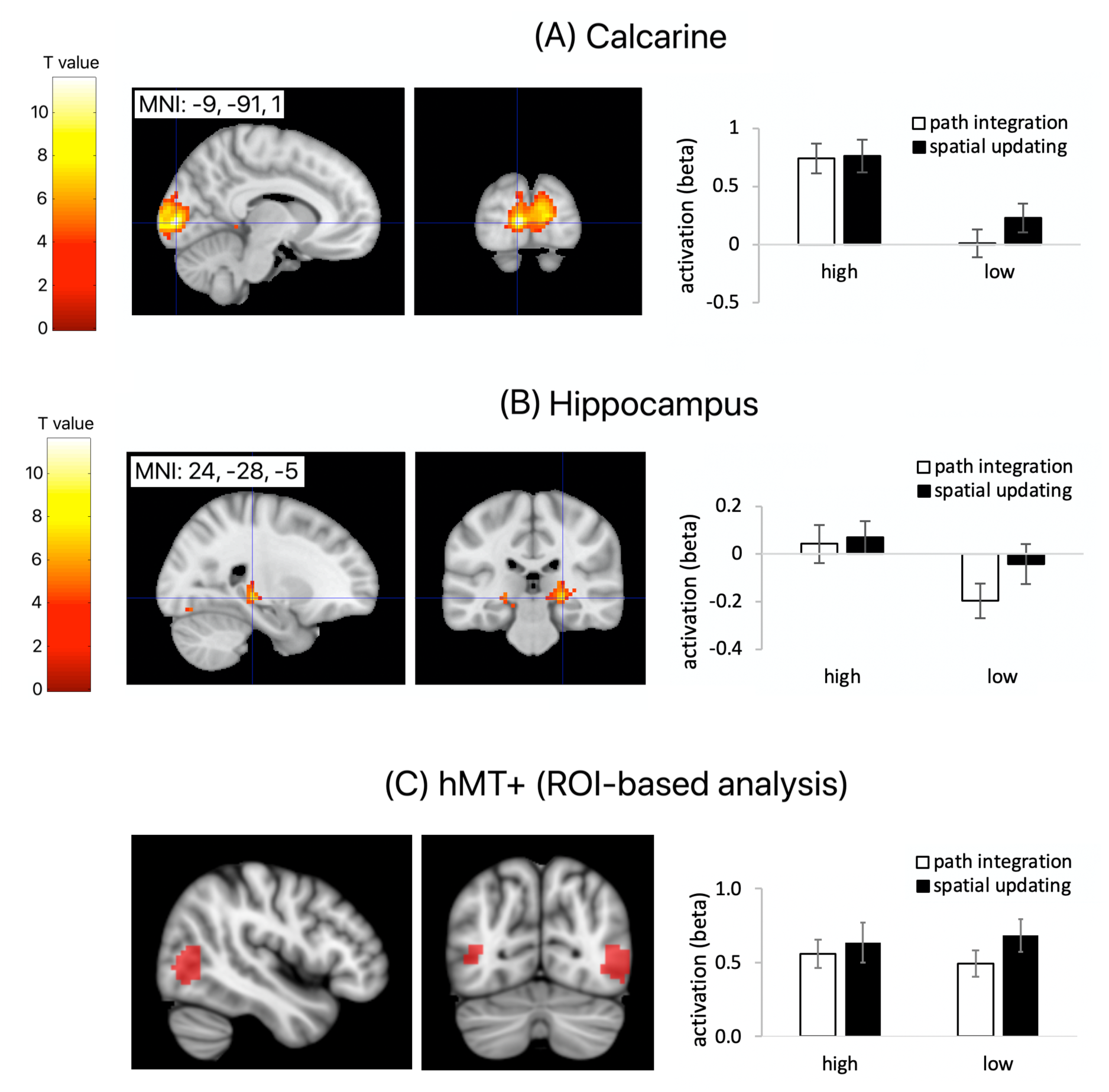
Main Effect of Dot Density on BOLD Responses in the Univariate FMRI Analysis. **(A)** V1 showed stronger activation for high- than low-dot-density trials, irrespective of task type. The left panel shows the statistical map projected onto the MNI template at a threshold of p_uncorrected_ < 0.001. The right panel plots mean activation estimates, averaged across voxels, within the significant cluster, as a function of task type and dot density. **(B)** Hippocampus showed stronger activation for high- than low-dot-density trials, irrespective of task type, following the same format as in (A). **(C)** hMT+ activation was not modulated by dot density. The left panel displays the anatomical hMT+ mask for one example participant. The right panel displays mean activation estimates (averaged across voxels) from a ROI-based analysis using participant-specific hMT+ masks, plotted as a function of task type and dot density. All other aspects are as in (A) and (B). Error bars represent SE of the mean.

**Table 2:**
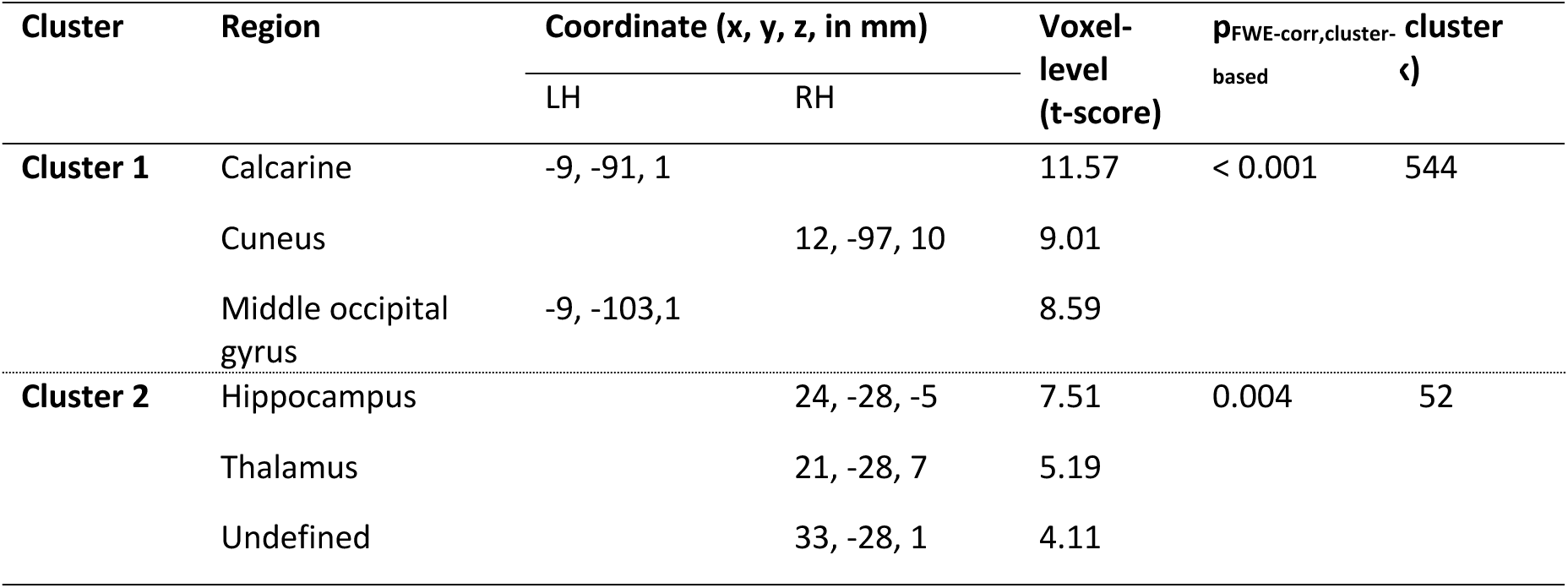
Main Effect of Dot Density in the Univariate fMRI Analysis. Spatial coordinates of the local maxima in the group analysis (p < 0.05 corrected). These regions showed higher activation for high vs. low dot density, independent of the task. Multiple comparisons were corrected across the entire brain, using the parametric cluster-based analysis (voxel-defining threshold: p_uncorrected_ < 0.001). Cluster size is determined with voxel-wise p_uncorrected_ < 0.001. RH/LH – right/left hemisphere.

Importantly, we did not find any significant voxels in areas known to subserve the analysis of optic flow. Specifically, despite identifying individual human motion complex (hMT+) with a separate functional localizer paradigm (see Methods), we found similar BOLD responses for both dot densities (Figure 7C) in this area (F(1, 15) = 0.02, p = 0.88, η_p_^2^ = 0.002). These results suggest that neuronal responses already reached saturation levels when only 750 dots were present.

#### 3.3.2. Psychophysiological Interaction (PPI) Analysis

The preceding univariate analysis revealed greater activation in spatial updating than path integration trials in the precuneus and the dorsal premotor cortex. To further investigate if path integration and spatial updating recruited dissociable neural mechanisms in other aspects of neural responses, we conducted the psychophysiological interaction (PPI) analysis, using the two areas as the seed regions. The PPI analysis examines whether effective connectivity from the seed region to other brain regions across the entire search volume differed between the two task contexts (Friston et al., 1997).

Recall that the behavioral results of this experiment could not arbitrate between the PI-depends-on-SU hypothesis and the PI-SU-independent hypothesis. A critical distinction between these hypotheses is that PI-depends-on-SU hypothesis predicts path integration relies on the output of spatial updating (Figure 1B), whereas the PI-SU-independent hypothesis posits the involvement of additional cognitive components different from the output of spatial updating, such as a head direction system that establishes a stable reference frame (Figure 1C). PPI is well-suited for testing this distinction, as it can reveal qualitative differences in effective connectivity between tasks, beyond the quantitative activation differences detected by the univariate analysis. If path integration requires computations beyond those of spatial updating, we would predict stronger connectivity between the precuneus or the premotor cortex with other brain regions, such as those supporting head direction processing.

Given the high sensitivity of connectivity analyses to outliers, we performed the PPI analyses using a first-level GLM in which outlier volumes were modelled as individual spike regressors. Outlier volumes were identified using the SPMUP toolbox (Pernet, 2021). Specifically, we calculated global signal intensity and framewise displacement for every volume. Volumes exhibiting atypical values on either metric were flagged as outliers using a robust S-estimator–based detection method (S-outliers), which provides high sensitivity to extreme observations while minimizing the influence of already-contaminated data. All other aspects of the first-level GLM were identical to those used in the mass univariate analysis.

When using the precuneus cluster as the seed region, we observed greater effective connectivity with the thalamus, superior frontal gyrus, and the inferior frontal gyrus in path integration trials compared to spatial updating trials (Table 3, Figure 8). These three clusters also reached statistical significance when a nonparametric permutation test was applied (thalamus, p_FWE-corr_ = 0.0196; superior frontal gyrus, p_FWE-corr_ = 0.0472; inferior frontal gyrus, p_FWE-corr_ = 0.0424). The opposite contrast (i.e., spatial updating > path integration) did not yield any significant voxels. These results suggest that while spatial updating activated the precuneus to a larger degree than path integration, the precuneus had stronger connectivity with the thalamus and the frontal areas during path integration.

**Figure 8.**
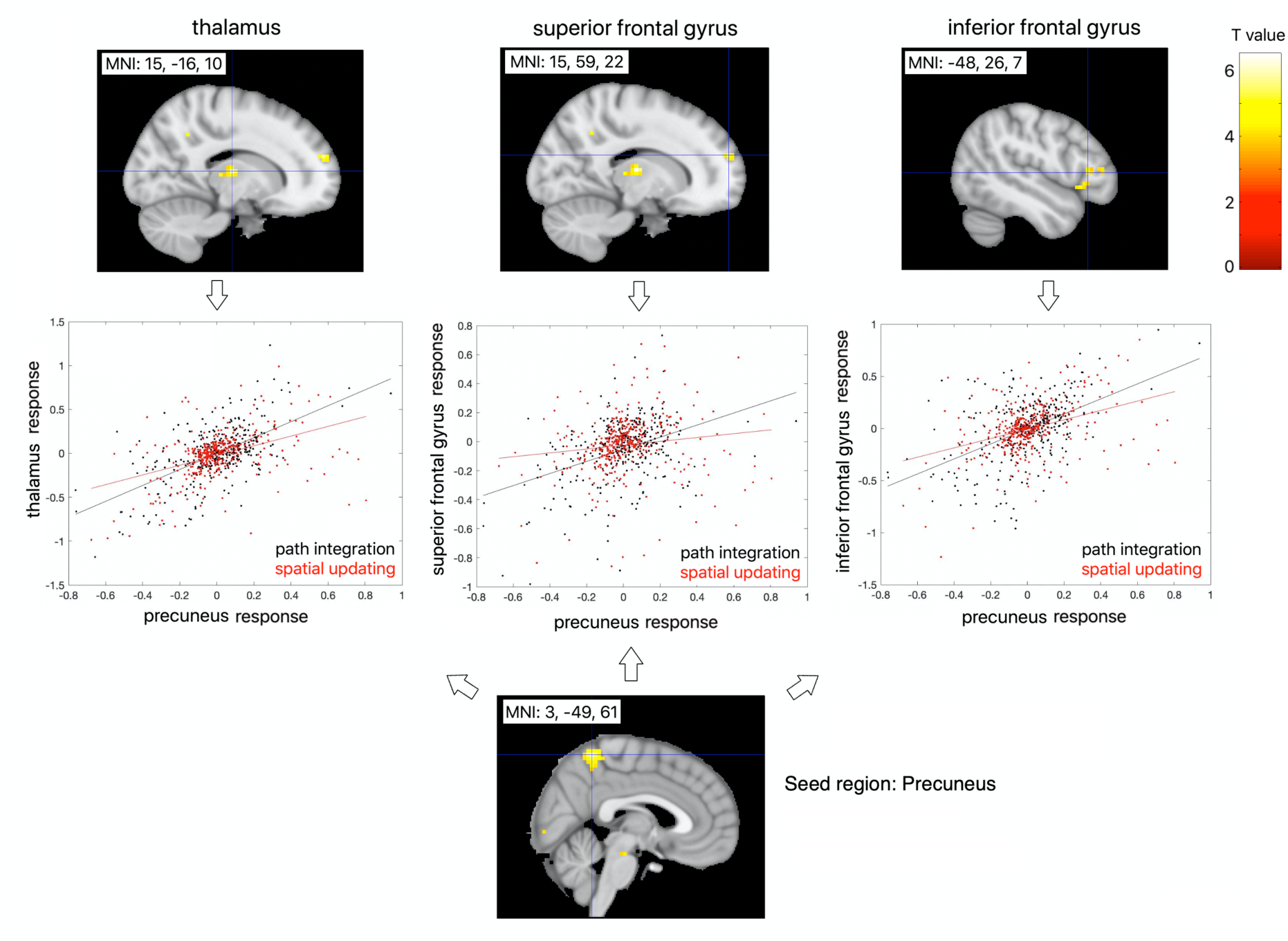
PPI Results with the Precuneus Cluster as the Seed Region. The seed region here corresponds to the precuneus cluster displayed in Figure 6A, which displayed stronger activation during spatial updating compared to path integration (lower row). In the upper row, PPI results are displayed on the MNI template at a threshold of p_uncorrected_ < 0.001. In the middle row, scatterplots depict the temporal relationship between brain activity between regions of interest for each task type, based on the first eigenvariates of the time-series BOLD signals for these clusters.

**Table 3.** PPI Results with the Precuneus Cluster as the Seed Region. Using the precuneus cluster (Figure 6A) as the seed region, three clusters of voxels showed stronger connectivity with the precuneus in path integration trials than spatial updating trials. The table shows the_spatial coordinates of the local maxima in the group analysis (p <. 05 corrected). Multiple comparisons are corrected across the entire brain, using the parametric cluster-based analysis (voxel-defining threshold: p_uncorrected_ < 0.001). Cluster size is determined with voxel-wise p_uncorrected_ < 0.001. RH/LH – right/left hemisphere.

| Cluster | Region | Coordinate (x, y, z, in mm) | | Voxel-level<br>(t-score) | $p_{\text{FWE-corr, cluster-based}}$ | Cluster size<br>(k) |
| --- | --- | --- | --- | --- | --- | --- |
|  |  | LH | RH |  |  |  |
| Cluster 1 | Thalamus |  | 15, -16, 10 | 6.48 | 0.004 | 50 |
|  |  |  | 6, -13, 4 | 4.73 |  |  |
|  |  |  | 15, -25, 7 | 4.72 |  |  |
| Cluster 2 | Superior frontal gyrus, dorsolateral |  | 15, 59, 22 | 5.45 | 0.033 | 32 |
|  | Superior frontal gyrus, medial |  | 6, 53, 22 | 5.32 |  |  |
| Cluster 3 | Inferior frontal gyrus, triangular part | -48, 26, 7 |  | 5.04 | 0.029 | 33 |
|  |  | -45, 35, 10 |  | 4.72 |  |  |
|  |  | -48, 20, -5 |  | 4.58 |  |  |
|  | Inferior frontal gyrus, orbital part |  |  |  |  |  |

When using the dorsal premotor cortex cluster as the seed region, we only detected a marginally significant cluster in the thalamus (peak voxel, MNI coordinate: [9, -13, 4]; cluster size = 27 at p_uncorrected_ < 0.001; _PFWE-corrected_ = 0.055), which tended to show stronger connectivity with the seed region in path integration trials compared to spatial updating trials. We obtained a similar result when applying a nonparametric permutation test (p_FWE-corr_ = 0.0658).

#### 3.3.3. Whole Brain Analysis of Connectivity Differences

The preceding PPI analysis showed that effective connectivity from a selected seed region differed between path integration and spatial updating, indicating that the two tasks recruit qualitatively different neural mechanisms. To examine connectivity differences at a broader, whole-brain level between these two processes, we examined inter-region functional connectivity using a multi-edge pattern analysis, based on 264 representative brain nodes (Power et al., 2011). For each task type, we computed a connectivity matrix capturing the pairwise correlations between all node pairs, and asked whether the pattern of these correlations across the entire brain was sufficiently different to reliably identify which task a participant was performing. As shown in Figure 9, a classifier trained on these connectivity patterns distinguished spatial updating from path integration with an accuracy of 84.375%, which was significantly higher than expected by chance (permutation test, p = 0.0002). This demonstrates that the two tasks induce reliably different large-scale network configurations, providing converging evidence that they engage distinct neural processes.

**Figure 9.**
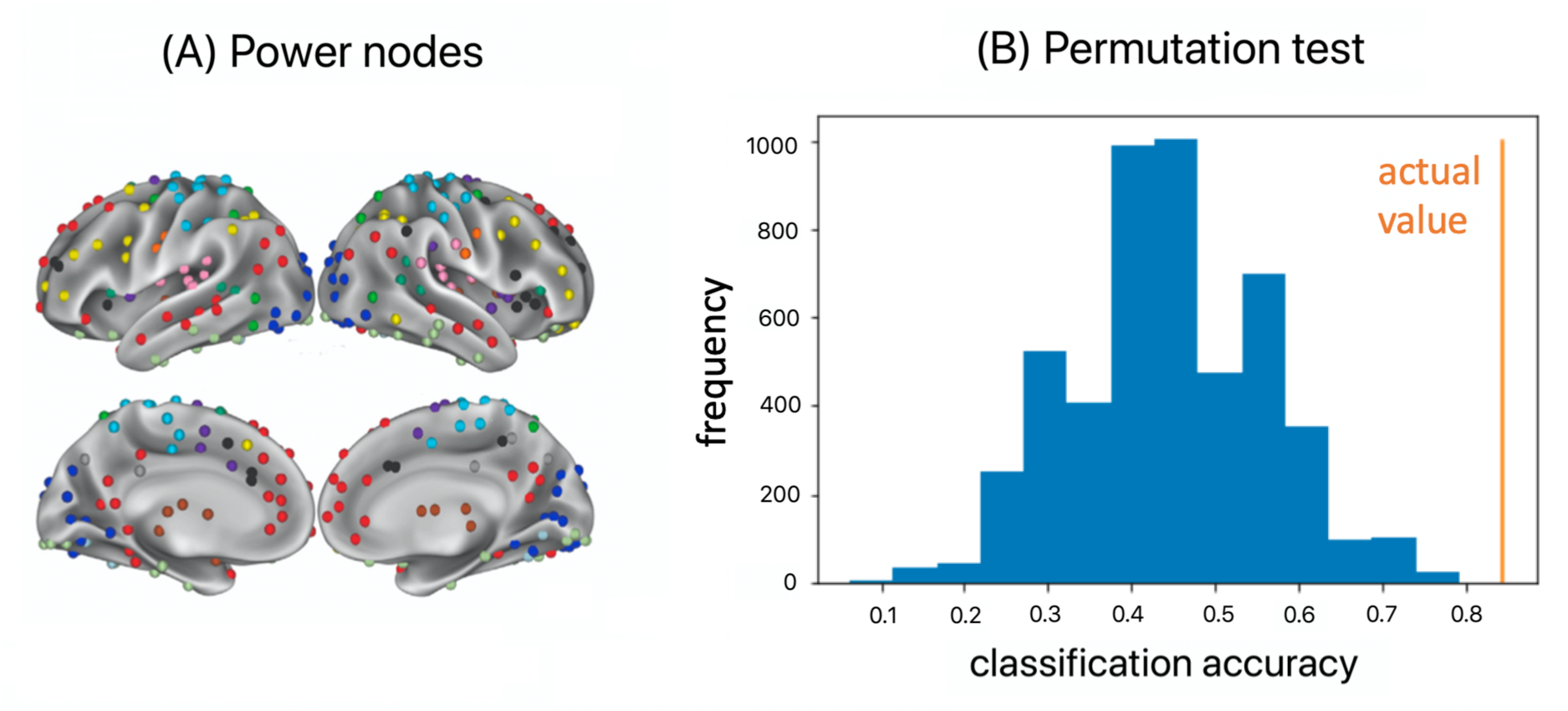
Results of the Whole-Brain Analysis of Inter-Regional Functional Connectivity Pattern. **(A)** Nodes used in the analysis, which are from Power et al. (2011). **(B)** Results of the inter-region functional connectivity pattern classification analysis. The blue bars represent the surrogate distribution of classification accuracy, which was obtained by randomizing the task labels of the trials. The orange bar represents the true classification accuracy (= 0.844), with a significance level of p_1-tailed_ = 0.0002.

#### 3.3.4. Summary of fMRI Results

The fMRI analyses provide additional evidence differentiating the three hypotheses. Across the univariate, PPI, and whole-brain multi-edge pattern analyses, path integration and spatial updating exhibited not only quantitative differences in activation but also qualitative differences in inter-regional connectivity.

In particular, although the precuneus showed greater activation during spatial updating, while its connectivity with multiple regions, including the thalamus, was stronger during path integration. These connectivity differences indicate that additional computations were recruited during path integration compared to spatial updating, contradicting the PI-depends-on-SU hypothesis and providing more direct evidence supporting the PI-SU-independent hypothesis. The thalamus has been shown to process head direction information anchored to the global environment in both rodents (Taube & Muller, 1998) and humans (Shine et al., 2016). One plausible interpretation is that salient directional information, such as the initial head direction, was used to establish a stable reference frame (He et al., 2018) within which the self-to-home vector could be computed (Figure 1C).

## 4. Discussion

In this study, we examined whether spatial updating and path integration - two navigation processes that rely on the integration of self-motion cues - are supported by shared or distinct cognitive-neural mechanisms. By isolating visual self-motion information in the form of optic flow, we directly compared the two processes under identical sensory input. Across behavioral, eye-tracking, and fMRI measures, our findings converge on the conclusion that spatial updating and path integration rely on relatively independent spatial computations. In the behavioral experiment, participants responded faster during spatial updating than during path integration, and eye-tracking revealed distinct task strategies associated with this reaction-time difference: while participants fixated the remembered target location during spatial updating, gaze was oriented in the direction of motion during path integration. Importantly, the reaction-time advantage for spatial updating persisted in the fMRI experiment even when eye movements were controlled, indicating that the behavioral dissociation cannot be attributed to overt differences in visual sampling. Critically, spatial updating was associated with stronger BOLD responses in the precuneus and dorsal premotor cortex, whereas path integration showed stronger task-dependent effective connectivity between the precuneus, thalamus, and frontal movement-related regions, pointing to a context-dependent reconfiguration of functional interactions. In addition, whole-brain connectivity patterns distinguished the two task states. Together, these findings support the view that spatial updating and path integration engage distinct cognitive-neural mechanisms, even when driven by the same visual self-motion information.

A central strength of the present study is the integration of behavioral testing, eye-tracking, and fMRI within a single experimental framework. This multimodal approach enables converging evidence across measurement modalities and helps offset the relatively small sample sizes typical of many fMRI studies. In particular, this approach allowed us to evaluate three competing hypotheses proposed in the literature (Figure 1A–C). Under the SU-depends-on-PI hypothesis, path integration is assumed to operate automatically, whereas spatial updating requires an additional inference step based on the output of the path integrator, resulting in worse behavioral performance (e.g., slower responses) for spatial updating. The PI-depends-on-SU hypothesis reverses this dependency: during spatial updating, participants continuously track the target location, while path integration requires additional computation based on the output of spatial updating at the response stage. This would predict worse behavioral performance (e.g., slower responses) for path integration due to additional computation, but attributes this cost to an inference process within a stable reference frame that is independent of spatial updating. Although the latter two hypotheses share the same behavioral prediction, they differ fundamentally in their proposed underlying mechanisms.

Behaviorally, while absolute pointing errors were comparable across the two tasks, responses were faster during spatial updating than during path integration. This pattern is incompatible with the SU-depends-on-PI hypothesis but remains consistent with both the PI-depends-on-SU and the PI–SU-independent hypotheses. Eye-tracking data provided critical additional constraints. During spatial updating, participants maintained fixation at the remembered target location until it left the field of view, despite the absence of a visible stimulus – a form of “looking at nothing” behavior that has been linked to goal-directed attention and facilitated memory processing in other cognitive domains (Ferreira et al., 2008; Henderson & Castelhano, 2005; Johansson & Johansson, 2014; Lakshminarasimhan et al., 2020). In contrast, during path integration trials, gaze was oriented in the direction of travel, a strategy associated with monitoring self-motion direction via optic flow (Authié et al., 2015; Authié & Mestre, 2012). Importantly, the reaction-time advantage for spatial updating persisted when participants had to maintain central fixation in the fMRI experiment, indicating that this effect cannot be explained by overt gaze behavior alone. Together, these findings suggest that both overt and covert spatial attention to the target location can facilitate processing during spatial updating, while path integration relies on a different computational strategy.

While these behavioral results were incompatible with the SU-depends-on-PI hypothesis they could not distinguish between the PI-depends-on-SU and PI–SU-independent accounts. Eye-tracking findings favored the PI–SU-independent hypothesis, yet the underlying cognitive processes remained unclear, motivating the fMRI experiment. A first key fMRI result was stronger task-related activation of the precuneus and the middle frontal gyrus (close to the dorsal premotor cortex) during spatial updating compared to path integration. This pattern is consistent with a core assumption shared by the PI-depends-on-SU and PI-SU-independent hypotheses, namely that the target location was continuously tracked during spatial updating, such that the pointing direction to the target is readily available at response. Consistent with the eye-tracking results, these fMRI findings support an egocentric updating strategy, in which the target’s location is represented and continuously updated relative to the observer’s changing position and orientation. Converging evidence from previous work supports a central role of the precuneus and the dorsal premotor cortex in egocentric updating of object locations during navigation. For example, activation in these regions scales with the number of tracked locations (Jahn et al., 2012; Wolbers et al., 2008), a defining feature of the egocentric updating (Wang et al., 2006). Moreover, the precuneus is causally involved in spatial updating, as TMS-induced disruption of this regions impairs participants’ behavioral performance (Müller et al., 2018). These activation-based fMRI results are therefore inconsistent with the SU-depends-on-PI hypothesis, which proposes a reversed dependency between the two navigation tasks. However, because both the PI-depends-on-SU and PI-SU-independent hypotheses predict continuous target updating, this activation difference in the precuneus and the dorsal premotor cortex alone cannot adjudicate between them.

The PI-depends-on-SU and PI-SU-independent hypotheses can be discriminated by the connectivity results. Although the precuneus showed stronger activation during spatial updating, its coupling with the thalamus was enhanced during path integration. The thalamus has been implicated in representing directional information relevant for navigation, including head-direction signals in rodents (Taube & Muller, 1998) and humans (Shine et al., 2016). Head direction signals in the thalamus can be anchored to the global environmental space (Shine et al., 2016). Under the PI-SU-independent hypothesis, stable heading information in the thalamus could serve to establish a fixed spatial reference frame during path integration trials, likely anchored to the participant’s facing direction at the beginning of a trial. At response, directional relations defined in this frame were computed to obtain an egocentric vector pointing to the starting position, which was represented in the precuneus and the dorsal premotor cortex. Such head direction computations based on the thalamus are not necessary for spatial updating in the present task. In contrast, the PI-depends-on-SU hypothesis does not naturally predict this task-specific engagement of thalamic connectivity, as it assumes that path integration operates primarily on the output of spatial updating, without requiring supplementary computations carried out in other brain regions.

Finally, whole-brain task classification analyses provided converging evidence for the PI-SU-independent hypothesis by showing that spatial updating and path integration can be reliably distinguished based on inter-regional functional connectivity patterns. These analyses captured large-scale interactions among distributed brain regions, defined using a widely adopted set of functionally meaningful brain nodes derived from meta-analytic and resting-state connectivity work (Power et al., 2011). The successful classification of the two task states indicates that spatial updating and path integration recruit distinct large-scale network configurations, extending beyond the focal precuneus–thalamus connectivity effects identified in the PPI analysis. Although this network-level dissociation is broadly consistent with the PI-SU-independent hypothesis, the specific computational mechanisms underlying these differences remain to be clarified.

Although the current study provides valuable insights into the cognitive-neural mechanisms underlying path integration and spatial updating, its limitations should be considered when generalizing the findings to other contexts. First, because participants could not physically walk during MRI scanning, the self-motion cues under investigation did not include body-based cues, which are an important source of spatial information in real-world navigation. Previous research has shown that during path integration, when body-based self-motion cues are available, participants can adopt an egocentric updating strategy by continuously tracking the starting location to themselves, provided that they are instructed to do so (He & McNamara, 2018).

Second, during spatial updating in the current study, participants were required to track the position of only a single target. With such a limited tracking demand, participants may have relied primarily on transient egocentric spatial representations maintained in working memory. However, real-life navigation in complex environments often requires tracking multiple objects. When the number of objects exceeded working memory capacity, individuals may instead rely on allocentric spatial representations stored in long-term memory, as simultaneously tracking of many objects is cognitively costly and ineffective. This tendency is especially pronounced when the objects form a regular layout (Mou et al., 2006). In this case, people may update their own position and only reconstruct object positions at the response stage, so path integration may serve as the basic mechanism; in these scenarios, the SU-depends-on-PI hypothesis may therefore be more plausible.

In addition to revealing distinct mechanisms for spatial updating and path integration, our results also inform how perceptual properties of visual self-motion input influence brain responses. Manipulating dot density strongly modulated activity in early visual cortex (V1/V2), with higher densities eliciting larger BOLD responses. In contrast, regions further along the visual motion and navigation pathways, including hMT+ and the precuneus, were insensitive to this manipulation. This pattern is consistent with findings from non-human primates showing that neurons in area MST, which are functionally related to human hMT+, are relatively insensitive to low-level visual features like dot density (Duffy & Wurtz, 1991). Because dot density provides detailed information about local visual structure but less information about movement trajectory or direction, these findings are consistent with a role of hMT+ and the precuneus in extracting higher-level self-motion information rather than encoding low-level stimulus properties.

Dot density also modulated responses in the hippocampus and thalamus, with stronger BOLD signals observed in the high-density condition. Although this effect was not a primary focus of the study, it may reflect indirect consequences of altered self-motion perception. Computational modeling suggests that the spatial sparsity of optic flow cues can bias perceived self-motion speed, thereby affecting path integration performance (Lakshminarasimhan et al., 2018), and intracranial recordings in humans show that hippocampal theta activity is modulated by movement speed during navigation (Goyal et al., 2020). Accordingly, variations in dot density may have influenced speed perception, which in turn modulated hippocampal and thalamic responses. However, in contrast to the neural findings, we did not observe reliable effects of dot density on behavior. This dissociation between brain activity and behavior may indicate that the dot-density manipulation was insufficiently strong to elicit measurable behavioral differences.

## 5. Conclusions

The present study examined whether spatial updating and path integration, while often treated as overlapping forms of navigation, engage distinct cognitive and neural mechanisms. Converging evidence from behavioral performance, eye movements, region-specific activation patterns, and connectivity patterns indicate that these processes are not interchangeable but instead rely on separate computations. This dissociation underscores the importance of considering spatial updating and path integration as complementary yet independent mechanisms in human navigation.

## Acknowledgements

We thank Masaki Miyanohara for help with stimulus programming and Jana Wendler for help with data collection. This work was supported by the European Commission (Marie Curie Outgoing International Fellowship awarded to T.W.), the Volkswagen Foundation (Fellowship awarded to J.W.), National Natural Science Foundation of China (NSFC #32100839, awarded to X.C.), STI 2030 – Major Projects (#2021ZD0200409), Cao Guangbiao High Science and Technology Foundation, Zhejiang University (ZJU 2020QN002, awarded to X.C.).

## Declaration of generative AI and AI-assisted technologies in the manuscript preparation process

During the preparation of this work the authors used ChatGPT in order to improve the language and readability of the manuscript, with caution. After using this tool, the authors reviewed and edited the content as needed and take full responsibility for the content of the publication.

